# Qualitative Determination of Bioactive Metabolites of Chyawanprash by Using Gas Chromatography-Mass Spectroscopy

**DOI:** 10.1101/2022.10.10.511551

**Authors:** K Rajesh Babu, T S M Saleem, V S Thiruvengada Rajan, M Madhu, S Chand Basha, U Narasimhulu

## Abstract

Chyawanprash is a rich source of all the major bioactive metabolites because of its authentic and natural composition and widely used due to its medicinal properties. In the present study, the precise flow of analytical methods was employed for the identification and characterization of unknown secondary metabolites in Chyawanprash. The phytochemical components of the whole formulation were extracted using n-hexane, ethyl acetate, and methanol by maceration which shows the presence of all the major secondary metabolite components through preliminary qualitative phytochemical screening. The GC-MS analysis of different extracts of Chyawanprash was investigated using Clarus 680 GC (Perkin-Elmer (U. S. A) 30 m X 0.25 mm ID X 250 μm DF) which revealed the presence of total 15 unknown secondary metabolites. Finally, the results showed that the study forms the basis for the phytochemical characterization of Chyawanprash extracts, the isolation of responsible bio-active metabolites and their biological activity is necessary for future studies.

## 1. Introduction

Medicinal plant life represents the eternal kindness of almighty for the continuance of life in the universe. The rise in the human population, inadequate provider of medication, the preventive value of treatments, facet effects of many allopathic medications and development of resistance to presently used drugs for infectious diseases has magnified the special emphasis on the utilization of traditional medicine within the mankind envisaged from the multi-varied products mainly as medicines for a wide sort of human ailments [1]. Plants produce a tremendous diversity of low molecular weight components called metabolites which continue to be widely practiced on several dispensations since ages and according to WHO the eighty percent of the world’s population depends on herbal medicine for their primary health care [2].

Chyawanprash was the oldest, most popular, and traditional Ayurvedic polyherbal formulation which is widely used as health promotive and disease preventive ‘Rasayana’ drug in India and elsewhere. Generally, it is potent antioxidant paste prepares through a synergistic blending of around 54 herbs and spices [3]. It is reported to have a rich source of vitamins, proteins, carbohydrates, dietary fibers, low-fat contents, and appreciable levels of major and minor trace elements such as Fe, Zn, Co, Cu, Ni, Pb, Mn, tannic acids and Carotenoids. It also provides several essential phytoconstituents namely flavonoids, alkaloids, Saponins, and phenolic compounds.

It’s been incontestable that the screening of natural products for bioactive compounds will be the fortunate approach for the invention of the latest drug leads. Thus the present study is designed to evaluate the phytochemical profile and gas chromatography-mass spectroscopy (GC-MS analysis of various extracts of Chyawanprash to reveals it’s medicinal and nutritional significance as well as for the identification and characterization of test materials.

## 2. Materials and Methods

### 2.1. Chemicals and reagent materials

All the solvents used for extracting the bioactive components from the whole formulation of Chyawanprash were purchased from Merck (Darmstadt, Germany) and all other chemicals and reagents used for phytochemical screening and GC-MS analysis were bought from Sigma-Aldrich (U.S.A)

The polyherbal formulation Chyawanprash was purchased from local Ayurvedic pharmacy and was tested for authenticity in the department of botany, Government Junior College, Rajampet, Andhra Pradesh, India 516115. The purchased samples were stored at 25°C and kept in their packaging until the time of analysis was performed within the expiry date.

### 2.2. Extraction of Bioactive components

The successive extraction of active biological components of the whole formulation with the help different solvents of increased polarity range was done through maceration by soaking the test material in n-hexane, ethyl acetate and methanol of rising polarity range one after the other in an airtight bottle for about 7 days with occasional shakings, later it was filtered through whatmann filter paper no.1 [4]. The solvents were removed by distillation using a rotary evaporator and the extracts were concentrated than dried in vacuum desiccators. The final concentrated extracts were further utilized for phytochemical screening and GC-MS analysis for the identification and characterization of bioactive metabolites with special emphasis on secondary metabolites.

### 2.3. Phyto-Chemical Profile of Chyawanprash [5, 6]

Qualitative phytochemical analysis was performed to assess the presence of phytochemical constituents in selective extracts of Chyawanprash. The preliminary screening of the individual sample extract for the confirmation of the presence of specific bioactive metabolites was done according to the methods described by CK Kokate, AP Purohit, SB Gokhale (Pharmacognosy 46^th^ edition, 2010) and SH Ansari (Essentials of Pharmacognosy 1^st^ edition, 2005).

A series of definite primary identification tests were performed individually for the all three extracted test material of the whole formulation of Chyawanprash categorically for alkaloids (Dragendroff’s test), glycosides (Baljet test), terpenoids (Liebermann-Burchard test), flavonoids (Lead acetate test), phenolic compounds (Ferric chloride test) and tannins (Gelatin test) and reported.

### 2.4. Analysis of Chyawanprash by GC-MS

The analysis of the selected fractions was performed using Clarus 680 (Perkin Elmer. U. S. A) GC-MS because GC interfaced to a mass spectrometer (GC-MS) method was a direct and fast analytical approach for the identification and characterization of biologically active principles of a complex organic and biochemical mixtures based on their relative abundance i.e. m/z ratio.

#### 2.4.1. Sample Preparation for GC-MS analysis

The 4ml of each sample extract of different solvents as mentioned above of 5mg/ml were extracted separately using a specific solvent of relative polarity to that of the test extract like Acetonitrile and methanol and then sonicate for 1 hour and kept aside for about twelve hours for GC-MS analysis. The final sample solution was then filtered through 0.25 μM PTFE filter and 2 ml of sample was injected at a split ration of 10:1 into the GC-MS instrument for analysis.

#### 2.4.2. Instrumentation specification for metabolomics by GC-MS

The GC-MS analysis of the effective extracts was performed using Clarus 680 GC (Perkin Elmer. U. S. A) and the GC interfaced to a mass spectrometer (GC-MS) equipped with Elite 5 MS fused silica capillary column packed with 5% biphenyl, ninety-five percentage of dimethylpolysiloxane [30m length X 0.25 mm film thickness X 250 μm df]. For GC-MS detection, an electron impact ionization energy system with ionization energy of 70 eV; was used. Helium gas (99.999%) was used as the carrier gas at a constant flow rate of 1ml/min and an injection volume of 1 μl was employed (split ratio 10:1) with injector temperature of 260°C during the chromatographic run and an ion source temperature of 230°C. The oven temperature was programmed from 60°C (isothermal for 2 min.) with a ramp of 10°C/min. up to 300°C and hold for 6 min. Mass spectra were taken at 70 eV; a scan-interval of 0.5 sec. with a scan range of 40-600 Da. The total GC running time was 32 min.

The relative percentage amount of each component was calculated by comparing its average peak area to that of total areas. Software adapted to handle mass spectra and chromatograms was Turbomass version 5.4.2 for the identification and characterization of unknown phyto-components.

#### 2.4.3. Data acquisition and processing

Interpretation of the GC-MS mass spectrum was conducted using the database of national institute standards and technology (NIST) 2008 having more than 62000 patterns. The spectrum of the unknown component was compared with the spectrum of the known components stored in the NIST library. The name, molecular weight, and structure of the components of sample extracts were ascertained.

## 3. Results and Discussion

The whole formulation of Chyawanprash was extracted with selective solvents of different polarity range as n-hexane, ethyl acetate, and methanol. Each extract was primarily analyzed qualitatively for preliminary phytochemical analysis which later confirms the unknown secondary metabolites using GC-MS. A total of 15 chemically diverse metabolites were identified of which 4-ethyl-2-hydroxy cyclopent-2-en-1-one in Methanolic extract, hexadecanoic acid, (3-Bromo prop-2-ynyl) ester in ethyl acetate extract and N-acetyl-2-ethoxy amphetamine of n-hexane extracts of Chyawanprash has significantly reported as major bio-active metabolites through GC-MS analysis.

### 3.1. Phyto-Chemical Profile

The preliminary phytochemical studies were important because the crude extracts possess a varied composition of bioactive components [7, 8]. The qualitative analysis of methanolic, ethyl acetate and, n-hexane extracts of a polyherbal formulation Chyawanprash revealed the presence of terpenoids, flavonoids, tannins, alkaloids, glycosides, and phenolic compounds. The presence of alkaloids, tannins and, phenolic compounds were found only in methanolic extracts while glycosides were observed in n-hexane extract whereas terpenoids were analyzed qualitatively in n-hexane and ethyl acetate extracts and flavonoids were present in Methanolic and ethyl acetate extracts (Table 1).

**Table 1:**
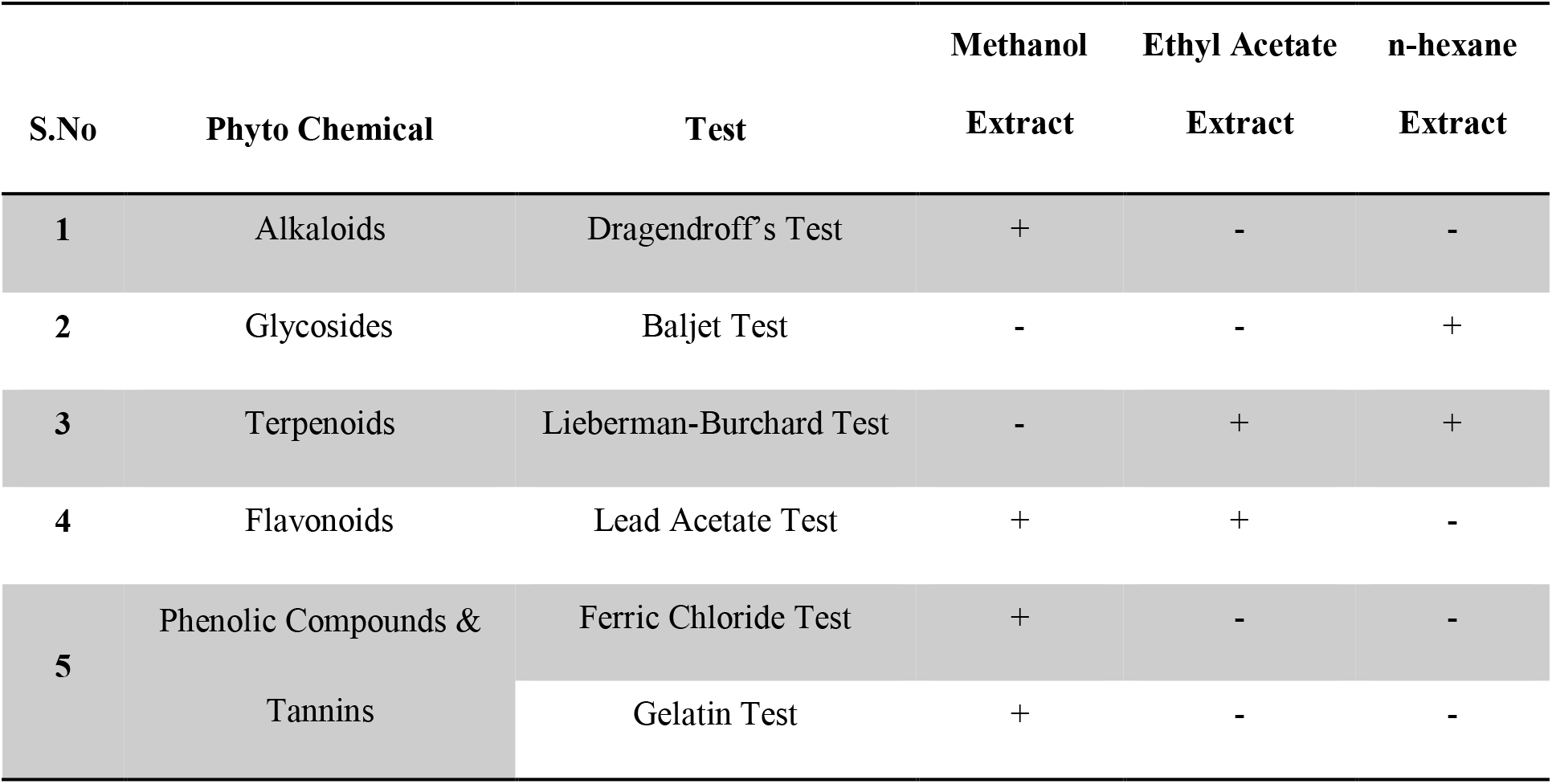
Preliminary Phytochemical screening of Chyawanprash.

Medicinal plants have therapeutic properties due to the presence of various complex chemical substances of different compositions which were found as secondary plant metabolites which contribute significantly to the new drug developmental studies from natural resources which have always been of huge interest to scientists working on infectious diseases [9].

The results expose that the formulation has quite a several different chemical constituents, which may be responsible for many pharmacological actions and have protective and disease preventive properties. The preliminary phytochemical profile of the polyherbal formulation, Chyawanprash gives an insight into its value as a medicinal as well as highly nutritious one, safe for consumption both as medicine and as a natural source as an immune booster.

### 3.2. Gas chromatography-Mass spectroscopy (GC-MS) analysis

However, the plant metabolomics not accepting the analysis of phytochemicals through GC-MS because of non-volatility and non-derivatization here we characterize major volatile metabolites of Chyawanprash using optimized chromatographic conditions. The reproducibility of the fragment mass spectral data of GC-MS was credible and fit well concerning to the NIST database [10]. The GC-MS chromatograms of different extracts of Chyawanprash showed majorly 15 metabolites which were putatively identified and characterized including 4-ethyl-2-hydroxy cyclopent-2-en-1-one, hexanoic acid, 3-hydroxy-ethyl ester, 3-propyl glutaric acid and 2-isopropyl-5-methyl cyclohexyl 3-(1-(4-chlorophenyl)-3-oxo butyl)-coumarin-4-yl carbonate in methanolic extract, hexadecanoic acid, (3-Bromo prop-2-ynyl) ester, octadecanal, 1-octadecyne, cyclohexane 1-(1,5-dimethyl hexyl)-4-(4-methyl pentyl)-, 4-pentadecyne, 15-chloro-, 1-hexyl-2-nitro cyclohexane in ethyl acetate extract, N-acetyl-2-ethoxy amphetamine, 1,3-bis-T-butyl peroxy-phthalan, 1-adamantine methylamine, α-methyl-, N-acetyl-3-methoxy amphetamine and 2,4,6-cycloheptatrien-1-one,3,5-bis-trimethylsilyl-in n-hexane extract (Table 2).

**Table 2:**
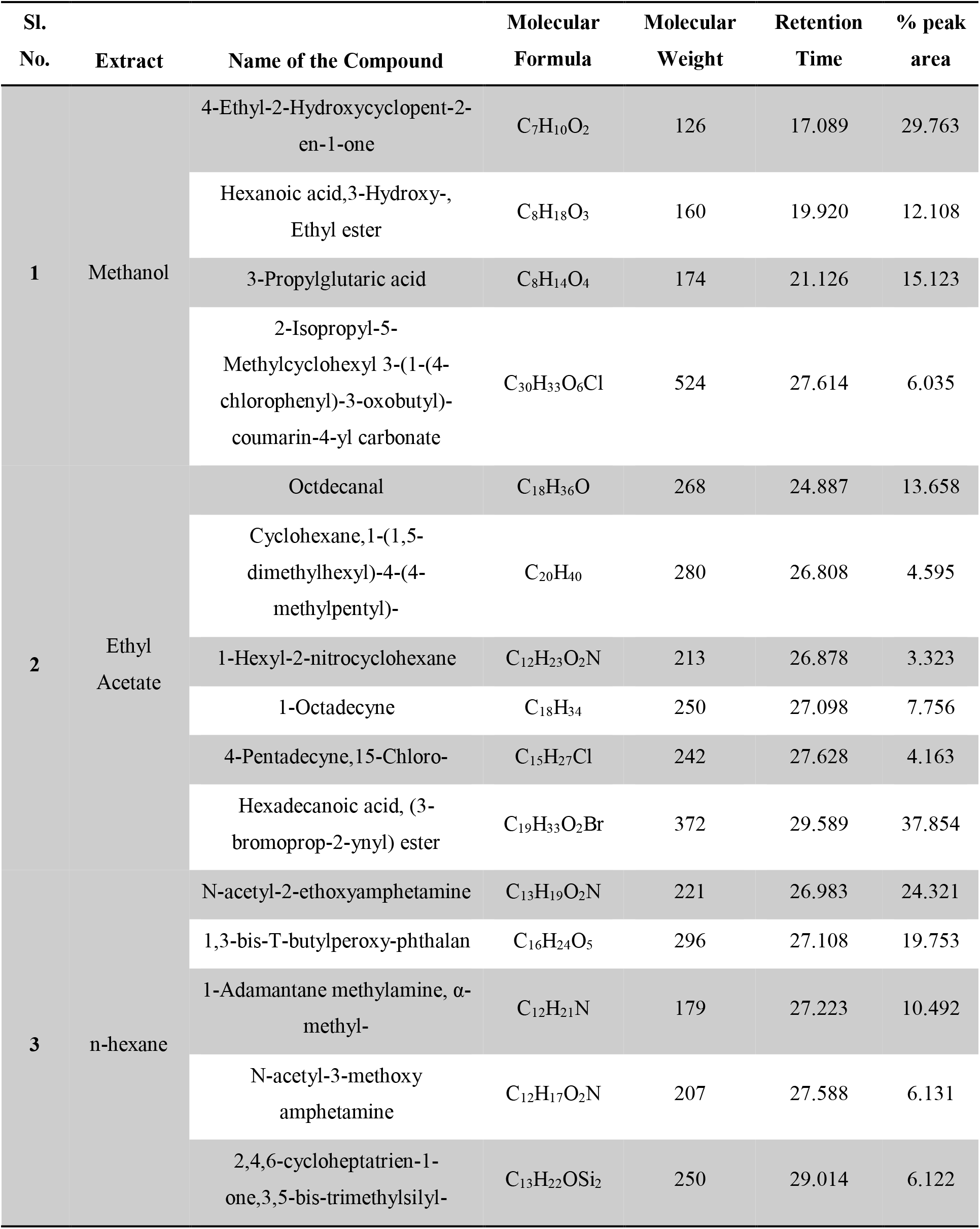
List of the bioactive metabolites identified in Chyawanprash by GC-MS.

The active principles with their retention time, molecular formula, molecular weight, and, % concentration are presented in table 2. The total ion chromatogram (TIC) of individual extract along with each fragmentation patterns of necessary compounds and the individual structures of identified bioactive metabolites were illustrated in fig. 1–3. Finally, the prevailing components with their contributing *in-silico* biological activity were presented by using the PASS online database in (table 3).

**Fig. 1.**
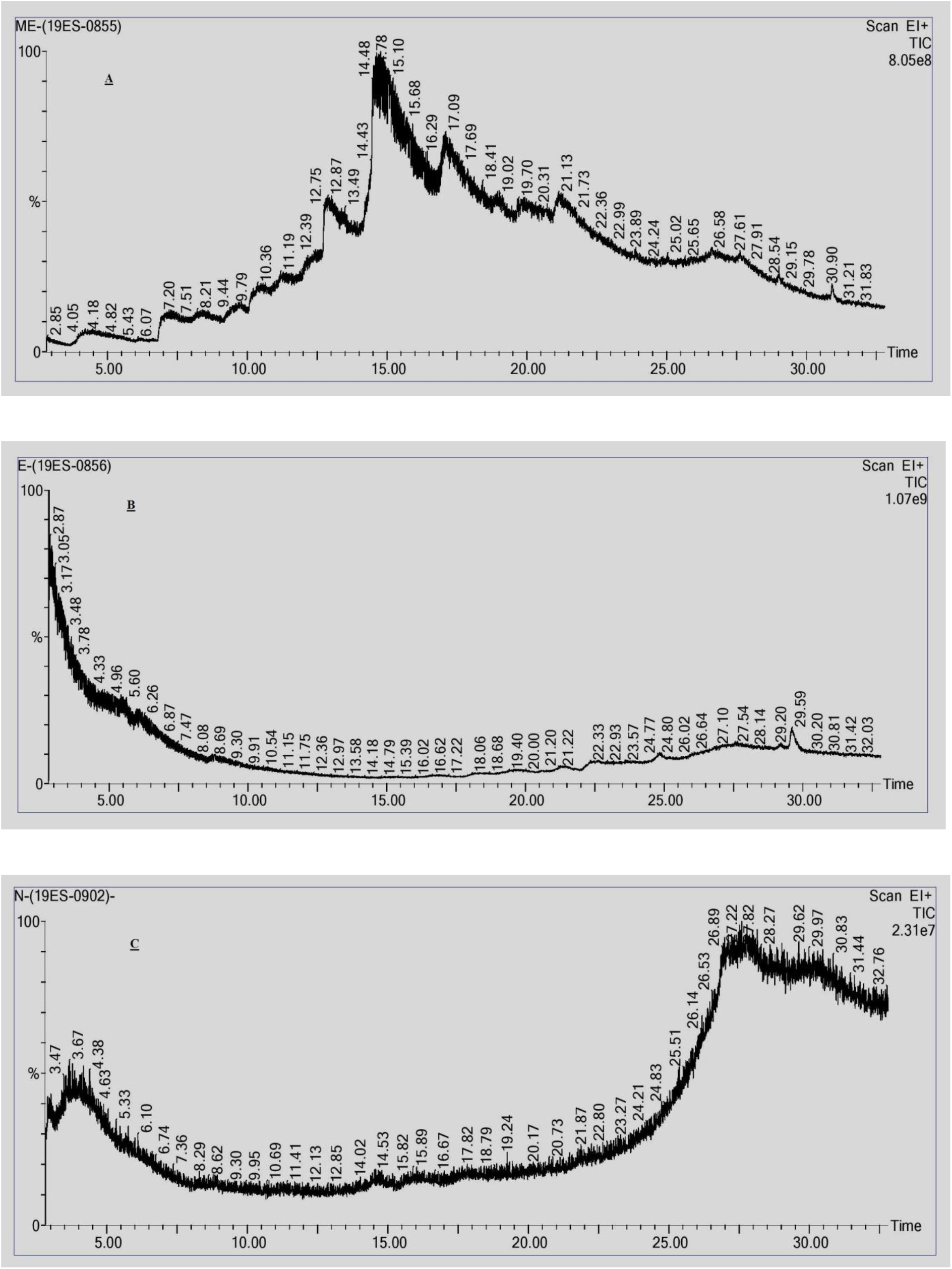
The GC-MS total ion chromatograms of differential extracts of Chyawanprash. (A) Methanolic Extract, (B) Ethyl Acetate Extract, (C) n-hexane Extract.

**Fig. 2.**
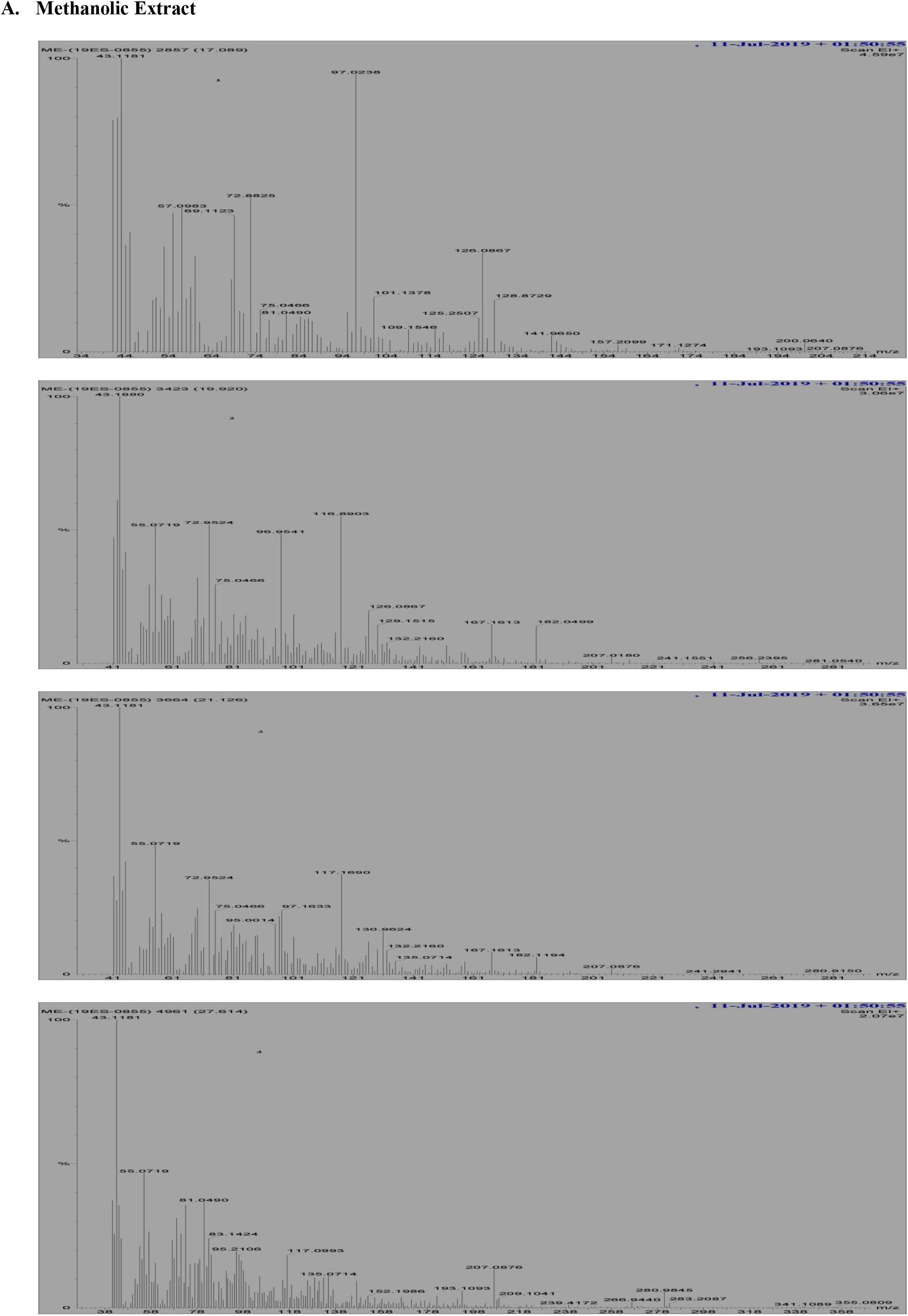
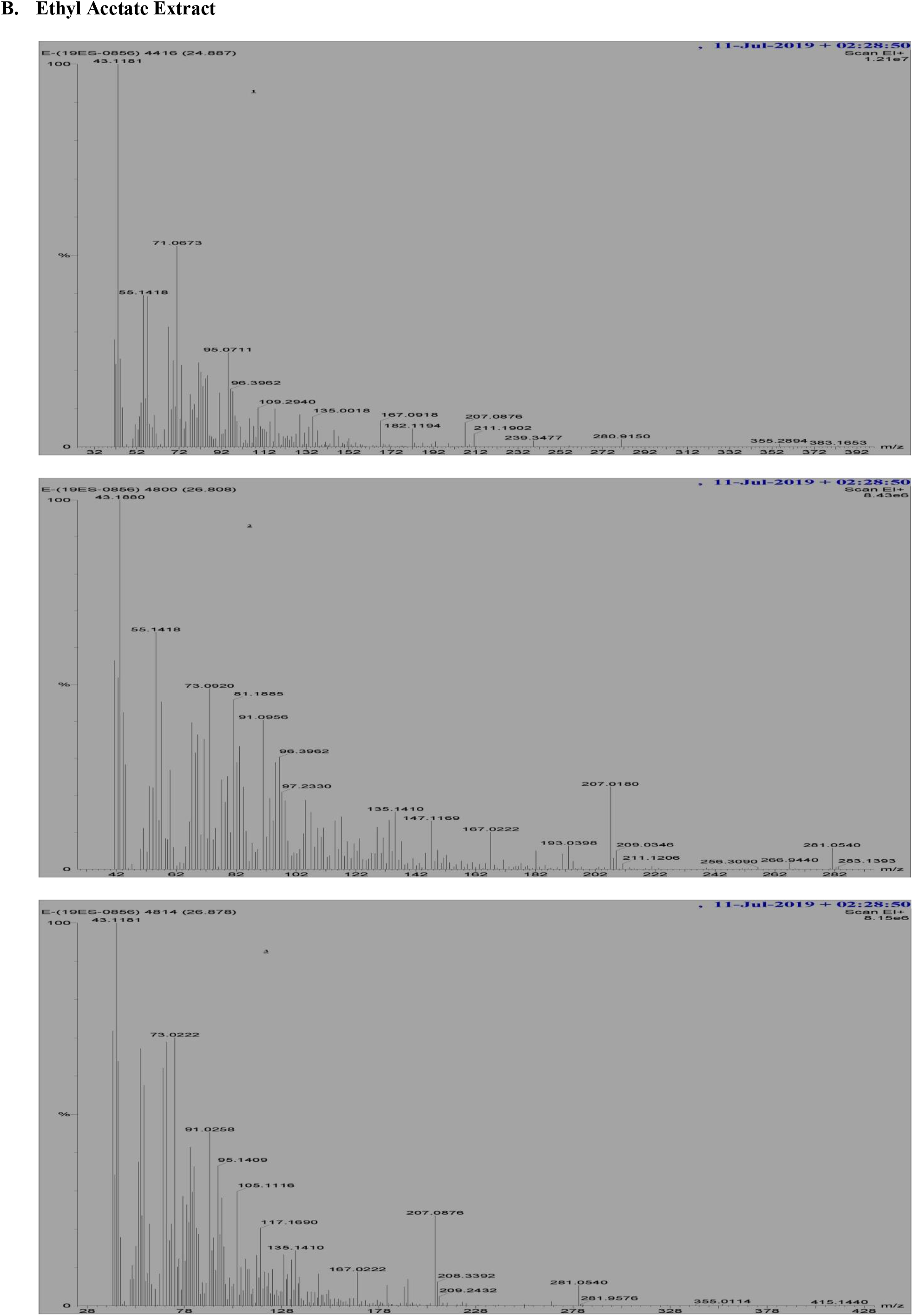
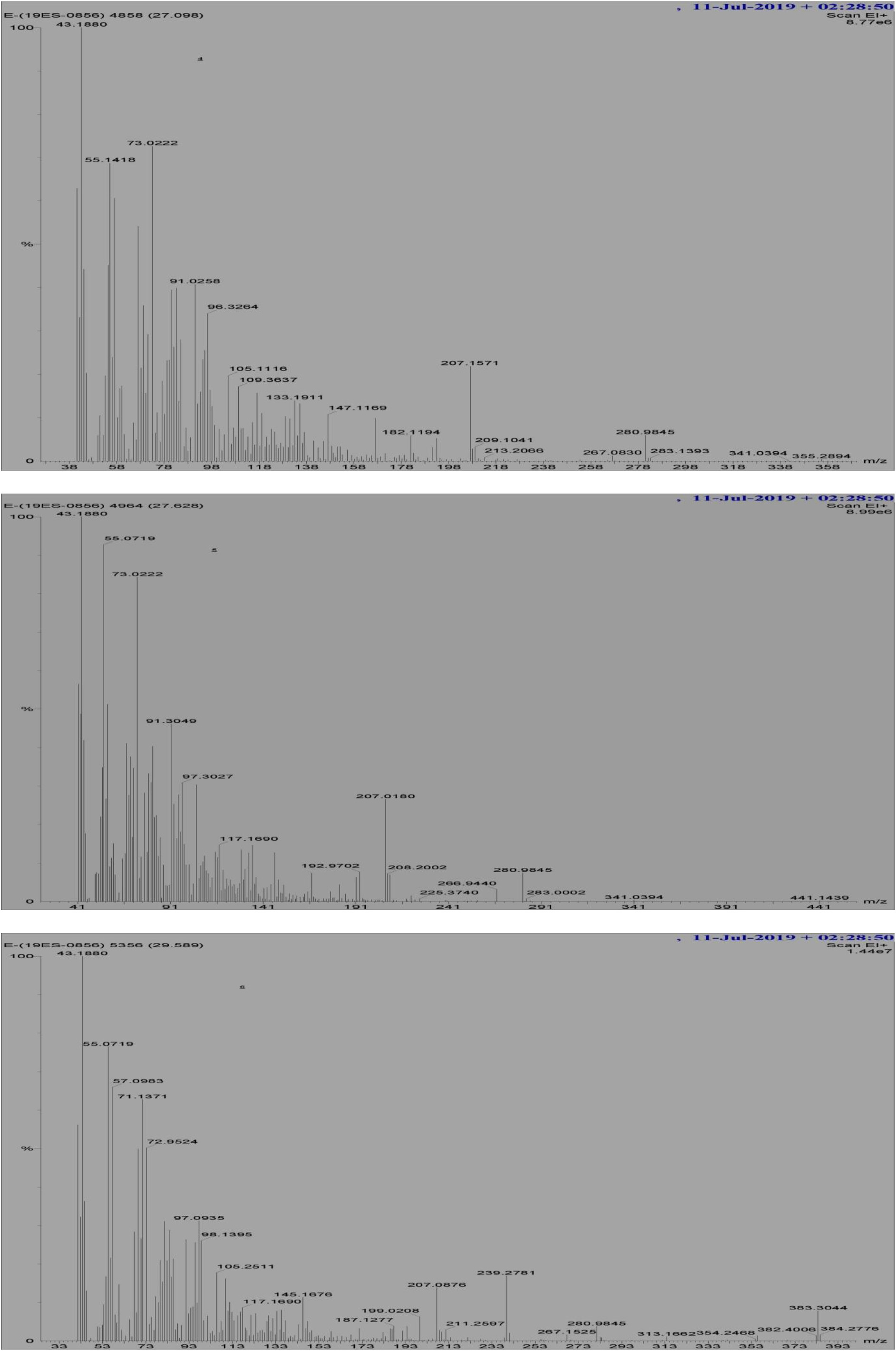
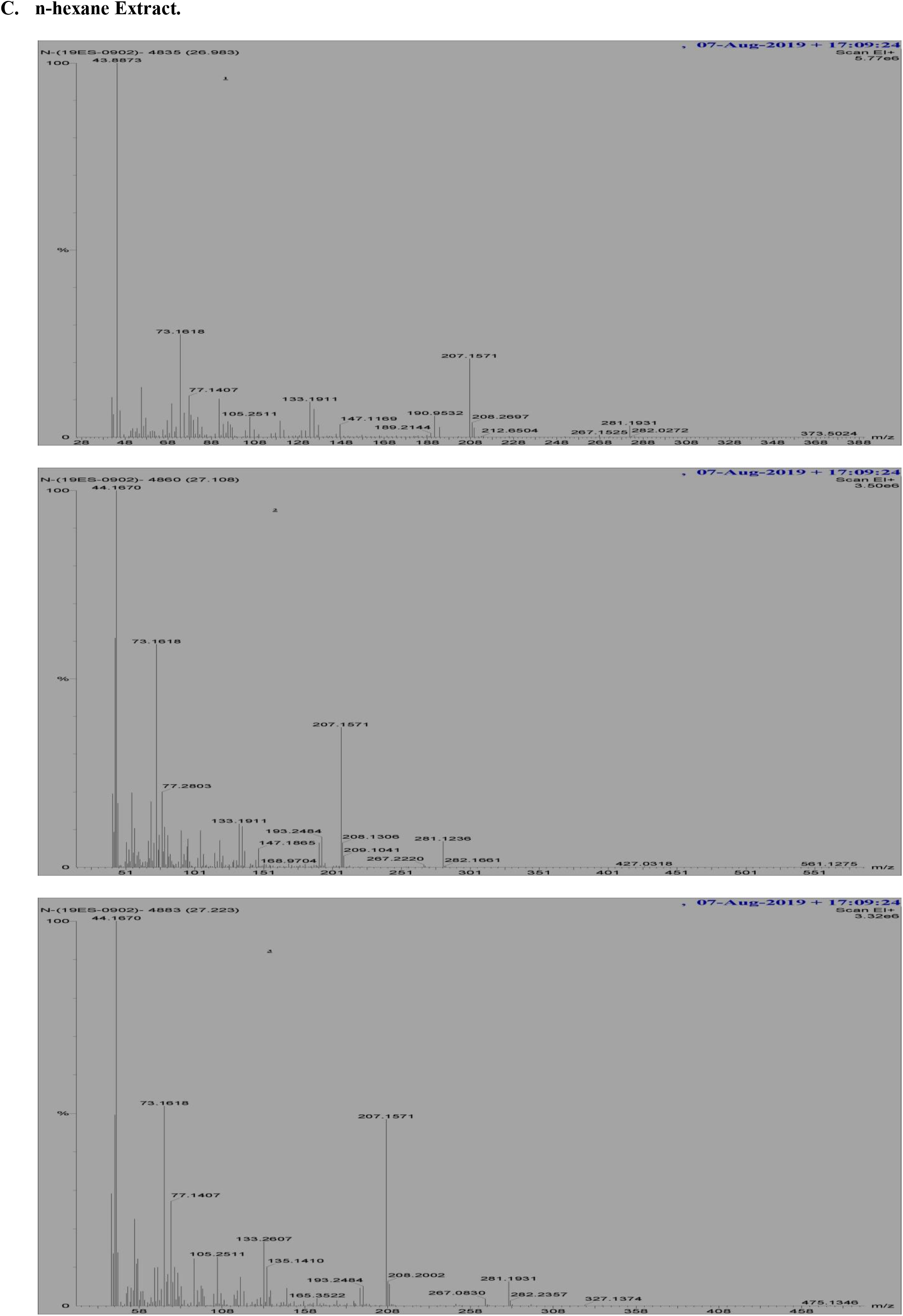
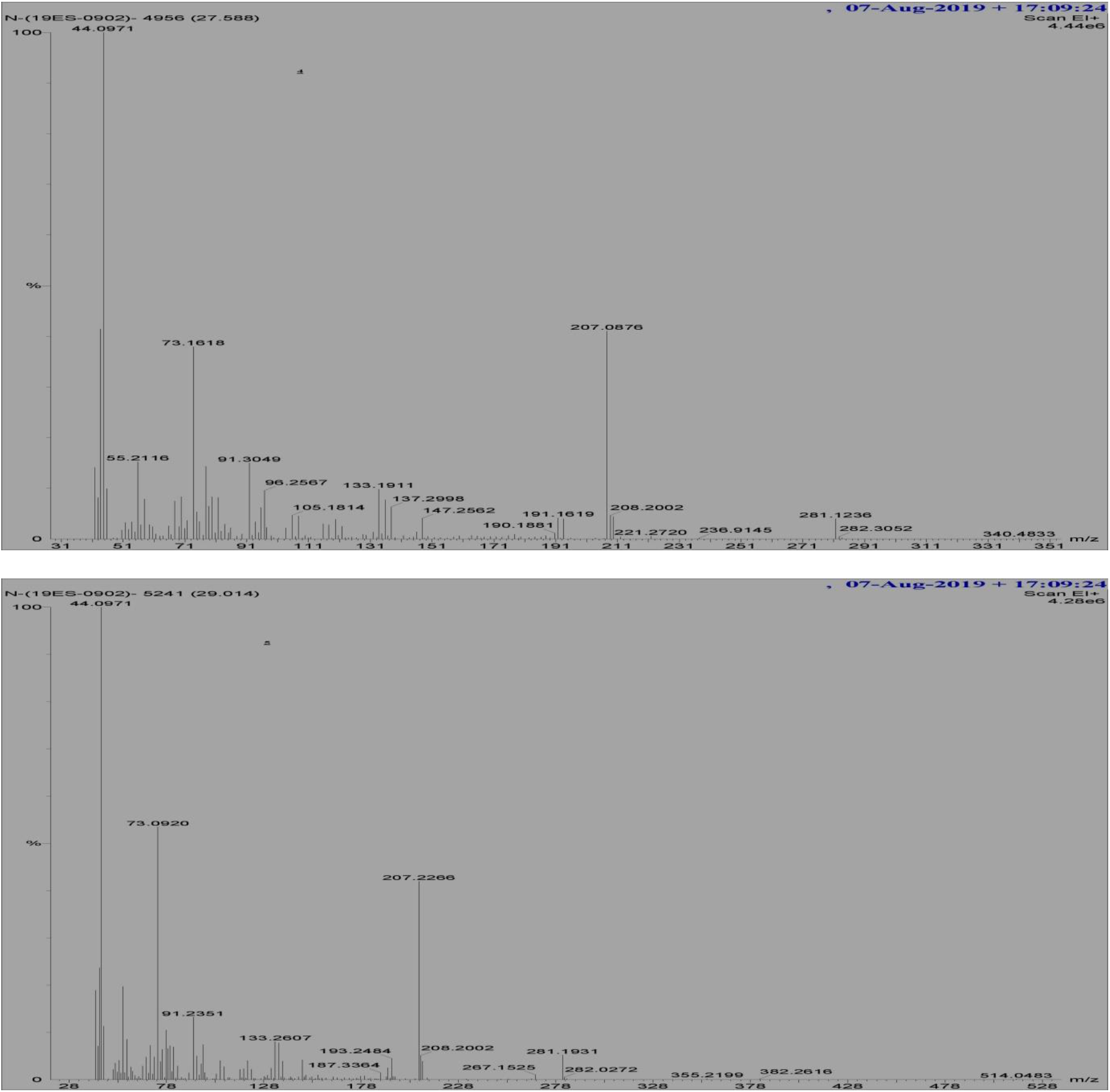
The GC-MS mass spectra of identified bioactive metabolites of (A) Methanolic Extract, (B) Ethyl Acetate Extract, (C) n-hexane Extract of Chyawanprash.

**Fig. 3.**
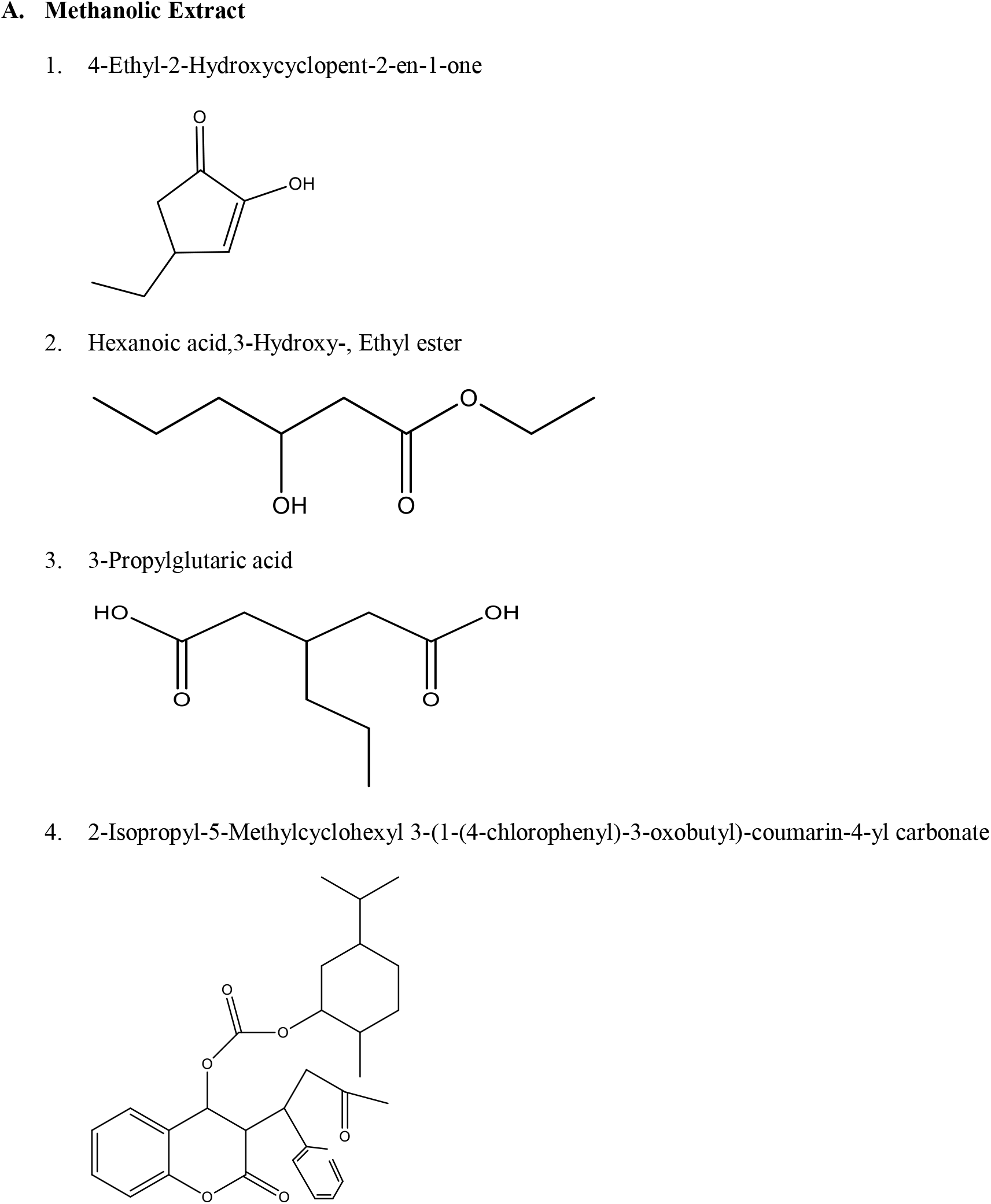
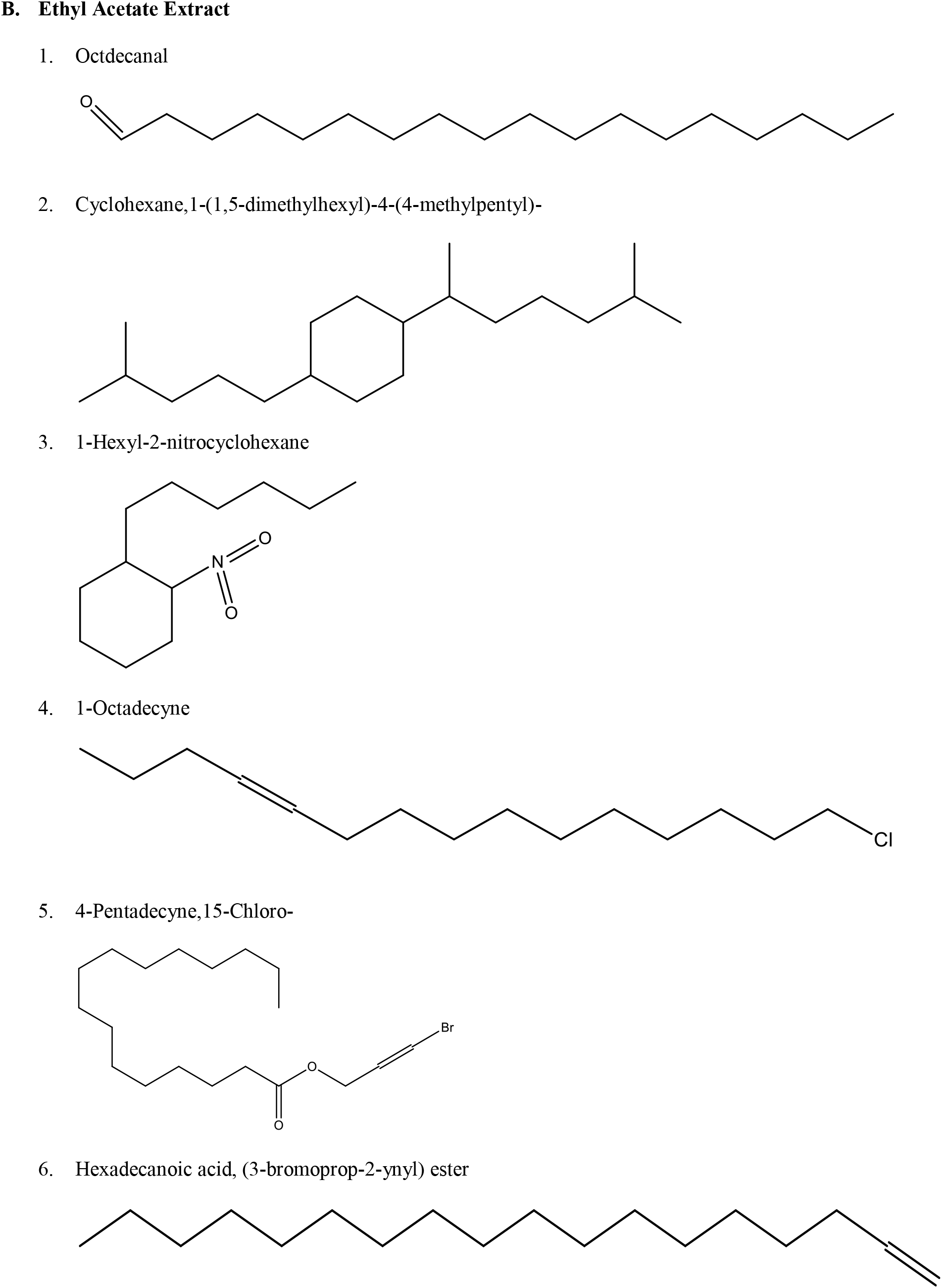
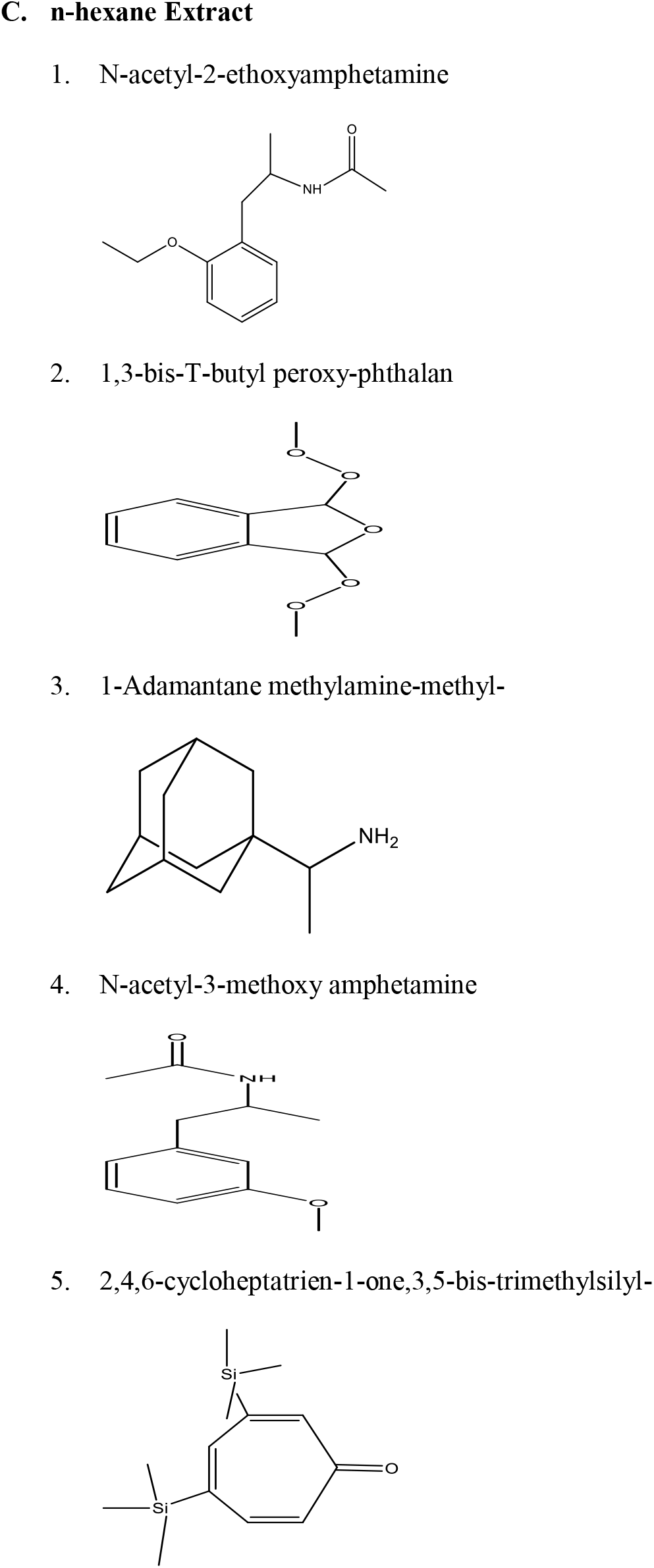
Structures of bioactive metabolites identified in differential extracts of Chyawanprash. (A) Methanolic Extract (1-4), (B) Ethyl Acetate Extract (1-6), (C) n-hexane Extract (1-5).

**Table 3:**
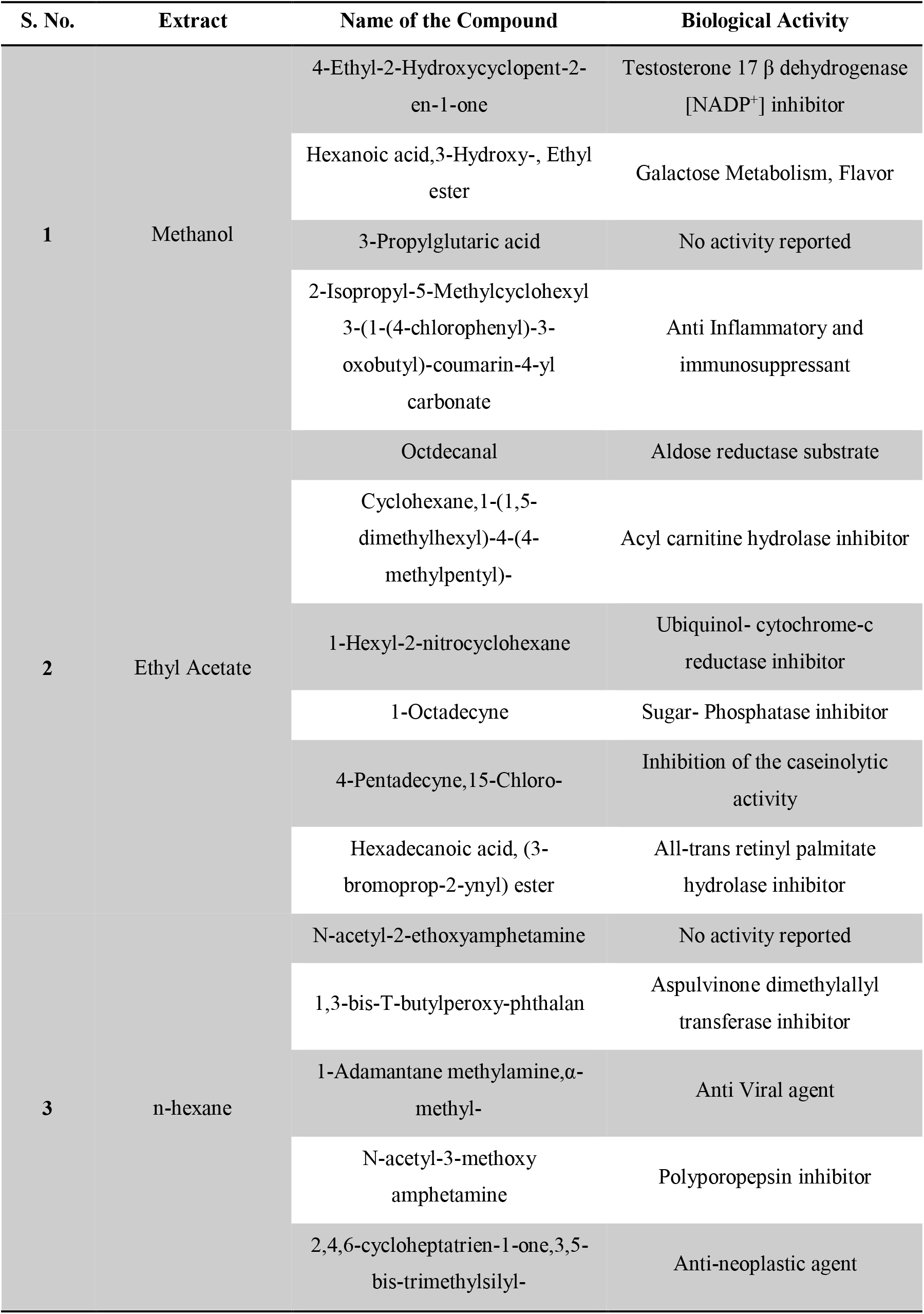
Biological Activities of phytochemical compounds identified in individual extracts of Chyawanprash.

The presence of these many bioactive metabolites in Chyawanprash justifies the utilization of the whole formulation for various ailments by traditional practitioners and also for new drug developmental studies for individual component analysis. A detailed study of the various compounds present in Chyawanprash and their pharmaceutical importance required to be carried such that a drug with multiple effects can be made available in the near future.

## 4. Conclusion

Although Chyawanprash was a comprehensive metabolic tonic with a blend of super concentrated nutrient-rich herb-mineral Ayurvedic recipe, the metabolomics analysis by GC-MS discriminates the origin and further provides a wide array of metabolites of the polyherbal formulation which includes organic acids, phenolic components, terpenoids, alkaloids which may act in major metabolic pathways with their significant actions. A total of 15 secondary metabolites was identified and characterized using GC-MS and all of these metabolites were identified by us for the first time in the whole formulation of Chyawanprash by using GC-MS. Isolation of individual phytochemical constituent and subjecting its biological activity may serve as candidates for new drug developmental studies in the treatment and prevention of many livestock diseases which furthermore has ample scope for future research and metabolic fingerprinting of herbal extracts is also essential for standardization of drugs which further establish the scientific basis for their pharmacological actions hence the study proves the versatile medicinal importance of Chyawanprash using gas chromatography-mass spectroscopy (GC-MS).

## Acknowledgments

Grateful acknowledgments are made to SIF-VIT Vellore, Tamilnadu, India, where all mass spectral studies were done. We are also thankful to Dr. T. Velpandian, Professor and In-charge of Ocular Pharmacology & Pharmacy, AIIMS, and New-Delhi for his assistance in technical design and analysis of research work.

## Abbreviations

GC-MS: Gas Chromatography-Mass Spectroscopy
NIST: National Institute of Standards and Technology
TIC: Total Ion Chromatogram
PTFE: Polytetrafluoroethylene (filter)

